# Evaluation of Probiotic Bacteria for the Reduction of Urea, p-Cresol, and Indole Levels

**DOI:** 10.1101/2025.03.24.644983

**Authors:** Sandra S. Hammer, Balendu Sankara Avvaru, Aditya Bahl, Steven C. Quay

## Abstract

**Introduction:** End-stage kidney disease (ESKD) is associated with the accumulation of uremic toxins such as urea, p-cresol, and indole, which significantly contribute to systemic complications such as inflammation, oxidative stress, and gut dysbiosis. Probiotics have demonstrated potential in modulating gut microbiota and reducing these toxins. We evaluated the *in vitro* efficacy of multiple probiotic strains of *Lactobacillus, Bifidobacterium, Sterptococcus, Bacillus* and others in degrading urea, p-cresol, and indole.

**Methods:** The probiotic strains were initially trained on toxin media prior to their evaluation of their toxin breakdown efficiency. Additionally microbiological methods using selective Christianson’s broth/agar and Stuart’s broth/agar were applied to assess urea breakdown. Liquid Chromatography-Mass Spectrometry (LC-MS) was utilized to quantify p-cresol and indole degradation.

**Results:** The results indicated that all probiotic strains exhibited significant activity in reducing urea, p-cresol, and indole concentrations in the culture media. In urea breakdown we observed an oscillation urea/ammonia ratio at 24 hr intervals, although a complete elimination of the ammonium byproduct was not feasible. Among all tested probiotics, *Lactobacillus* species showed the highest efficiency in urea breakdown. Furthermore, the toxin removing efficacy of the probiotics was evaluated in a simulated gut environment using the TNO invitro gut model.

**Conclusion:** Cumulatively, the findings suggest that probiotics could offer a promising strategy for the reduction of uremic toxins and their associated complications in ESKD patients.

## Introduction

Uremic syndromes are considered to be associated with uremic toxins and are compounds that accumulate in patients with end-stage kidney disease (ESKD)^1,2^. ESKD is characterized by a progressive decline in renal function, leading to the systemic accumulation of uremic toxins such as urea, p-cresol, and indole in systemic circulation. Urea is a nitrogenous waste product, while p-cresol and indole are protein-bound solutes derived from microbial fermentation of dietary tyrosine and tryptophan, respectively. Research thus far indicated that uremic toxins, namely indoxyl sulfate or p-cresyl sulfate, also increase the risk of cardiovascular disease or even mortality in patients with chronic kidney disease^5^. Elevated levels of these toxins contribute to inflammation, cardiovascular complications, and increased mortality in CKD patients. Various disorders in patients with ESKD are associated with uremic toxins, and it is difficult to remove some of these toxins through dialysis^3,4^. These toxic compounds are not only involved in the etiology of uremic syndrome but also have the potential to cause cardiovascular disease and renal osteodystrophy.

The gut microbiota plays a key role in the production and metabolism of these toxins^6^. In the United States, over 20 million individuals suffer from kidney-related disorders, with more than 600,000 experiencing kidney failure^7^. Patients with ESKD often exhibit disruptions in gut microbiota^6,8,9^. These include elevated levels of urea and ammonia in the gut, compromised intestinal barrier integrity, and systemic inflammation. This gut microbiome plays a critical role in immune development and function. Dysbiosis in ESKD patients often results in enhanced microbial generation of p-cresol and indole, which are further converted to their toxic derivatives, p-cresyl sulfate and indoxyl sulfate in the liver^10^. Hemodialysis effectively removes small water-soluble solutes but struggles to eliminate protein-bound toxins such as p-cresol and indoxyl sulfate. The accumulation of these toxins is linked to complications like cardiovascular disease, infections, and higher mortality rates in dialysis patients^4^.

Probiotics have been suggested to improve the condition of ESKD patients as they can restore microbial balance in the intestines, delivering positive health effects when administered in the correct dosages^11^. However, the ability of specific probiotic strains to metabolize uremic toxins remains poorly understood^12^. For probiotics to be effective, the microorganisms must survive exposure to gastric acid and bile, colonize the intestinal lining, and compete against harmful pathogens. Recent studies also suggested that probiotics, particularly species of Lactobacillus, Bifidobacterium, and Bacillus, may help modulate the gut microbiome, thereby reducing the production and accumulation of uremic toxins. For ESKD patients, probiotics offer a complementary therapy option, particularly for those undergoing dialysis or awaiting kidney transplants^7,11,13,14^.

We studied probiotic strains capable of efficiently degrading uremic toxins and explored their potential as adjunct therapies for ESKD patients. We have further tested the probiotics in the TIM-2 system, which is an *in vitro* model developed to replicate the human digestive system, from the mouth to the large intestine^15^. The system consists of different compartments that simulate the stomach, small intestine, and colon, each of which can be controlled and monitored independently. This system allows for a high degree of accuracy in replicating the processes that occur during digestion and absorption, providing an ideal environment to test the effects of probiotics. The results obtained from the TIM system also corroborate that performed with conventional microbiological methods and support the utility of using probiotics for ameliorating the burden of uremic toxins in ESKD patients.

## Methods

### Probiotic strains

The following ATCC strains were employed in the study: *Bacillus pasteurii* (11859), *Bacillus coagulans* (7050), *Erwinia herbicola* (33243), *Streptococcus faecium* (BAA-2320), *Streptococcus thermophilus* (19258), *Bifidobacterium longum* (15707), *Bifidobacterium breve* (15700), *Bifidobacterium infantis* (15697), *Lactobacillus casei* (15008), *Lactobacillus bulgaricus* (11842), *Lactobacillus rhamnosus* (21052), *Lactobacillus plantarum* (BAA-793), *Lactobacillus gasseri* (33323) *Lactobacillus acidophilus* (BAA-2845) and *Akkermansia muciniphila* (BAA-835). All probiotic strains were grown on gradient plates and subsequently transferred to broth conditions under the recommended ATCC conditions. (see supplementary methods for further details).

### Urea/Ammonia Determination

Urea breakdown assay was performed in 96-well plates to measure total urea degradation in media and quantify the ammonia released using the Nessler’s reagent method by measuring absorbance at 420 nm (see supplementary methods for further details).

### LCMS measurement of p-cresol and indole

The quantification of p-cresol and indole in test samples was performed using liquid chromatography-mass spectrometry. The samples were extracted using acetonitrile. Calibration curves for indole (0.4–400 µM) and p-cresol (2–2000 µM) were prepared prior to the sample run, and QC samples were analyzed twice, before and after sample injections, with all results falling within the linear range. Samples with concentrations exceeding the calibration range were diluted appropriately. Chromatographic analysis was performed using a Shimadzu Nexera X2 HPLC system coupled to a SCIEX Triple Quad 4500 mass spectrometer with Analyst 1.7.0 software. Separation was achieved using a Kinetex C18 column (100 × 3.0 mm, 5 µm), and ionization was conducted using a heated nebulizer source. (see supplementary methods for further details).

### Invitro Gut Model

The TNO invitro gut model (TIM-2 system) was employed in this study, which is elaborated in detail elsewhere^16^. The TIM-2 System was inoculated with a mixture of equal ratios of probiotic strains with a cumulative 10 billion CFUs in 125 ml in simulated ileal effluent medium(SIEM) and dialysis solution. The system was kept under strict anaerobic conditions. The main study included two runs with urea and one control run without urea where each run lasted 24 hours. At the start of the adaptation period, the System was inoculated with approximately 35 ml of thawed probiotic mixture and 90 ml dialysis fluid and was allowed to adapt to SIEM for 16 hours at pH 5.5. After the adaptation period, filter-sterilized urea was added to the colonic lumen to reach a concentration of 25mM (1.5 g/L), mimicking the levels found in renal patients the urea gradient was maintained through slow addition against luminal dialysis. For the metabolite analysis, dialysate and lumen samples were collected at: t = 0, 8 and 24 hours after the start of the test period and analysed for urea and ammonia (see supplementary methods for further details).

## Results

Probiotic bacteria have gained significant attention for their potential applications in addressing environmental and health-related challenges. Among these, training probiotic strains to degrade specific toxins holds promise for managing conditions associated with toxin accumulation, such as end-stage kidney disease. This study aimed to isolate high-efficiency strains from each species by training them on a gradient plate containing increasing concentrations of urea, p-cresol, and indole—toxins that accumulate in the bloodstream of ESKD patients due to impaired renal function. Probiotic strains were exposed to this gradient to evaluate their adaptive capabilities. The gradient plate was prepared by solidifying a layer of agar infused with high concentrations of the toxins, followed by a second layer containing decreasing concentrations, enabling gradual exposure to simulate selection pressure. (Fig 1). Colonies that grew closer to the high-toxin end of the plate were isolated for further analysis.

**Fig 1.**
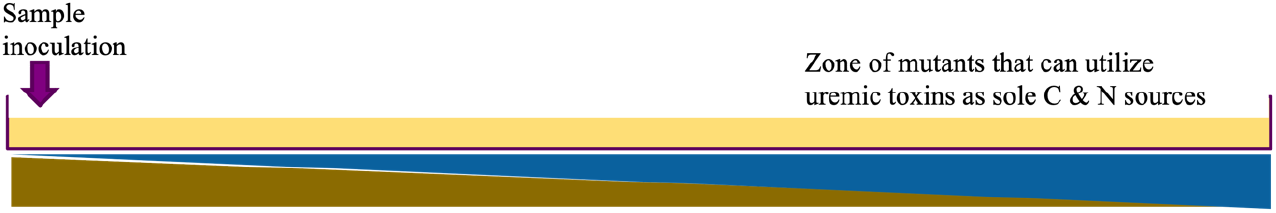
Illustration of a gradient agar plate used to assess the effects of an increasing toxin concentration (urea, indole or p-cresol) on microbial growth. The plate was made with a continuous toxin gradient, with the lowest concentration on the left side and the highest concentration on the right side. The microbial inoculum was evenly spread across the plate, and after incubation, distinct growth patterns emerged. For all strains on the left (low toxin concentration), robust microbial growth is observed, indicated by a dense, confluent lawn of colonies. As the toxin concentration increases toward the right, a gradual reduction in growth is visible, with a zone of partial inhibition followed by a complete absence of growth at the highest toxin concentrations. For each toxin, the right-most colonies were isolated for downstream experiments.

Results revealed strain-specific adaptations. To varying degrees, all the tested strains exhibited moderate growth predominantly in mid-gradient regions, and the high-efficiency isolates demonstrated significant degradation of urea, a key contributor to uremic toxicity. Plenty of colonies were formed in low-toxicity zones, but only a few colonies were visible on the gradient plate as the toxin concentration increased. The colonies formed in the zones of high toxin concentration were extracted and further cultured in liquid broth maintaining selection pressure of the toxins, and glycerol stocks were prepared for subsequent experiments. Among the tested probiotics, *Lactobacillus species* showed robust growth even at high-toxin concentrations.

### Urea breakdown in liquid media

Following the isolation of high-efficiency probiotic strains capable of degrading urea, p-cresol, and indole on gradient plates, the next phase of the study focused on evaluating their toxin breakdown capabilities in liquid media. All probiotic strains demonstrated significant urea-degrading activity compared to controls. Each of these probiotic strains were cultured in liquid culture media with 50% dilution to recommended media by ATCC for each strain. These culture media were spiked with urea concentrations between 50 to 300 mM. In the urea range tested, the results show all strains were able to break down urea efficiently. Notably, urea degradation exhibited an oscillatory pattern, with alternating high and low degradation rates observed at 24-hour intervals (Fig 2). For example, urea degradation peaked on days 1, 3, and 5, followed by reduced activity on days 2, 4, and 6. This cyclical pattern suggests potential regulatory mechanisms or metabolic shifts in the probiotic strains over time that trigger higher consumption of urea and ammoniums ions. However it was interesting to note that the corresponding OD values at 600 nm were relatively stable indicating a constant cell density

**Fig 2.**
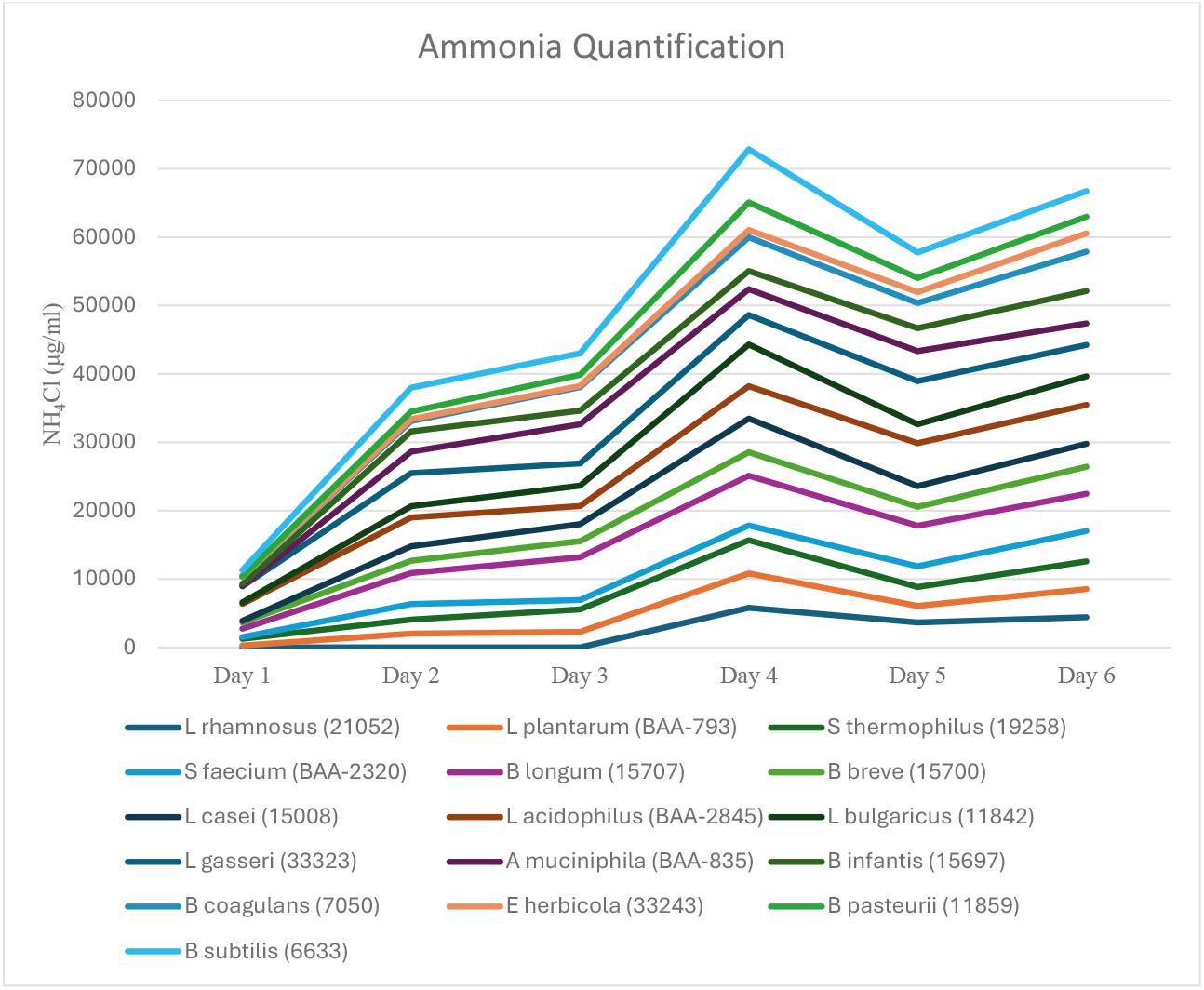
Growth and urea metabolizing characteristics of probiotics. Plot ammonium ion concentration in the media over a period of 6 days as measured using Nessler’s reagent. The plot is show as a stacked line chart for clearly discerning the NH4Cl oscillation trend during the experiment, where the Y-axis indicates cumulative ammonia concentration in the stack. Each line corresponds to a specific bacterial strain, where the stacking method visually represents trends in ammonia oscillation while maintaining comparative differences between strains. The raw data of the absolute values for the stacked line chart are given in Table S1 along with corresponding cell densities at OD600 in Table S2. The observed oscillation of ammonium ion concentration is likely a starvation response that triggers mechanisms for uptake of ammonium ions.

### Indole and p-Cresol Degradation

The degradation of indole varied significantly across strains and incubation times. The probiotic strains were incubated in minimal media supplemented with 5mM indole and assessed for their indole degradation capacity at the end of week 1 and 2 through LC-MS analysis. The LC-MS data confirmed a time-dependent decrease in toxin concentrations (*p* < 0.05). For indole, the control group showed concentrations of 386 µM and 421 µM (with 20% dextrose), while the test group showed 258 µM and 216 µM, corresponding to reductions of 127.5 µM in MM1 and 205 µM in MM1 + 20% dextrose conditions. Among the tested strains, *Lactobacillus species* demonstrated the highest efficiency, reducing indole concentrations 90% by the end of week 1 (Fig 3). Other species belonging to *Bifidobacterium, Bacillus* and *Streptococcus* genus showed moderate consumption indole with a reduction of 40 – 60% within a two week period. *E. herbicola* was the least efficient strain that displayed indole degradation showing a modest reduction of ∼20% after two weeks of incubation (Fig 4). In contrast to indole, almost all of the tested probiotic strains were only able metabolize p-Cresol to∼30% from 0 to 4 weeks of incubation. The only exception was *E. herbicola* which was able to metabolize >60% of p-cresol in the minimal media at the end of two weeks. It is interesting to note that unlike the rest of the probiotic strains that were able to source their carbon source from indole, *E. herbicola* which was unbale to breakdown indole could successfully metabolize p-cresol. Although both indole and p-cresol are aromatic compounds, their respective catabolic mechanisms seem to differ in the probiotic strains.

**Fig 3.**
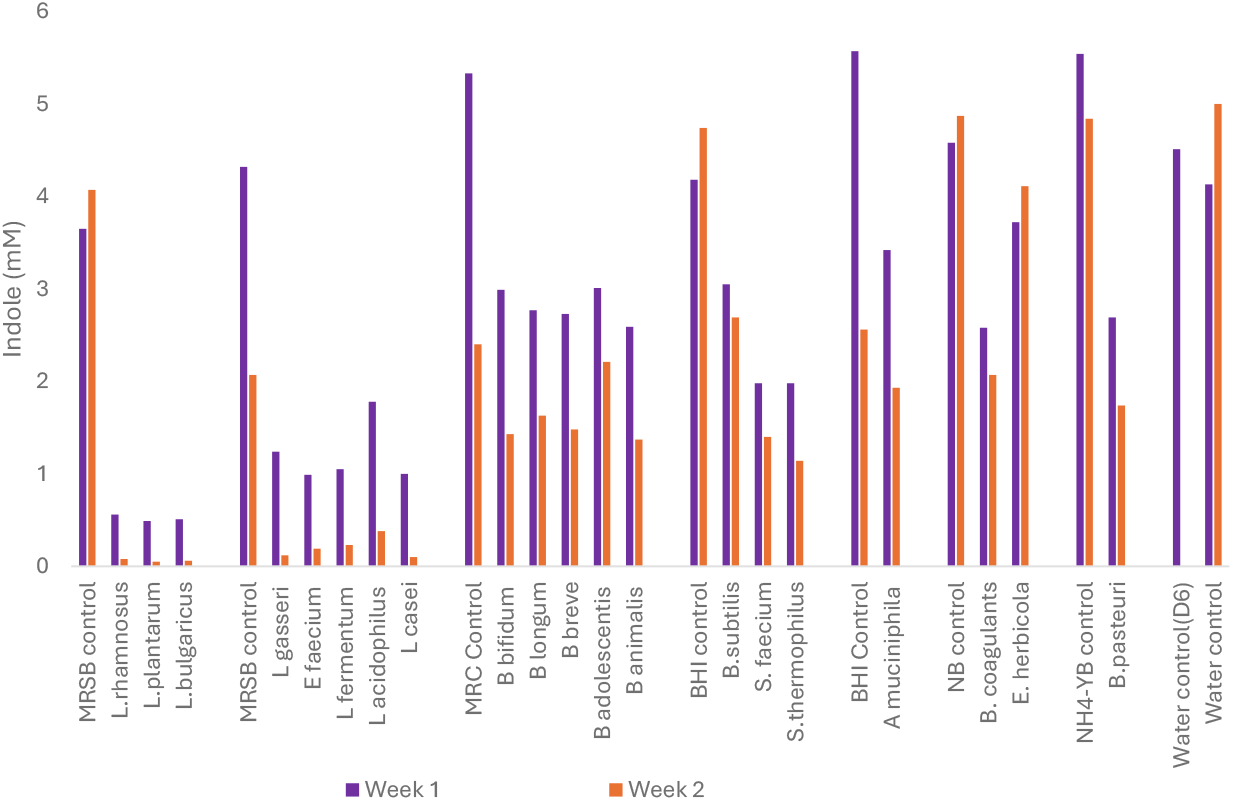
Consumption of indole by probiotics. The consumption of indole by the probiotics was followed for two weeks with a starting concentration of 5 mM. At the end of week 1 and 2, small quantities of the individual cultures were spun down and the supernatants were analysed on the mass spectrometry for determining indole concentrations.

**Fig 4.**
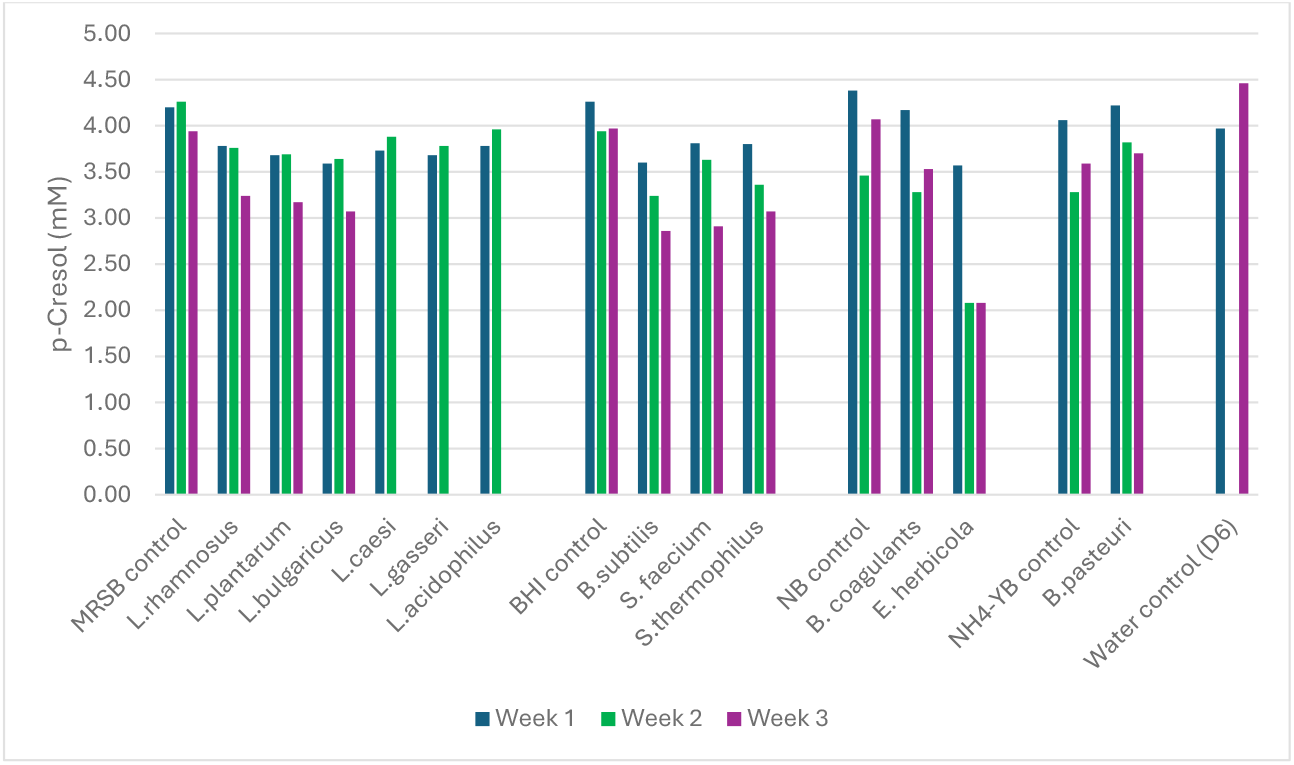
Consumption of p-Cresol by probiotics. The consumption of p-Cresol by the probiotics was followed for two weeks with a starting concentration of 5 mM. At the end of week 1 and 2, small quantities of the individual cultures were spun down and the supernatants were analysed on the mass spectrometry for determining p-Cresol concentrations.

### Invitro Colon Model

The probiotic mixture previously trained on urea metabolism was evaluated in the ‘invitro colon-model’ system to assess urea breakdown and further consumption of the ammonia byproduct. The invitro colon-model was made up of two main compartments of lumen and dialysate separated by a permeable membrane for metabolites but not the probiotics, which were restricted to the lumen compartment. The experiment was performed twice (exp. A and B) with a starting CFU count of 10^10 bacteria per run under a continuous supply of minimal media (dextrose and salts) supplemented with urea (25mM) at flow rate of 1.5 ml/min into the lumen. The samples were collected from the lumen and dialysate compartments at time points of 0, 8 and 24 hours for analysis of urea and ammonia concentration. A single control run was performed under identical conditions without urea supplementation.

The production of free ammonia was determined in the dialysate and lumen samples and are presented in Table S3-S6. The ammonia production (as measured in lumen plus dialysate) over the 24 hour time interval was 10.7 mmol. The urea concentration and amounts are shown in Table S4. These were corrected for the addition of urea over time. The urea consumption by the probiotic bacteria over 24 hours was 7.5 mmol. The settings from the pilot phase were also applied during the main runs. Only adjustment was the bead beating of the lumen samples to release urea from the bacterial cells. The probiotic cell counts were similar between experiments (i.e. with and without urea) and compared to the pilot phase. The results indicated that a differential gradient of urea was observed between the lumen and dialysate compartments between time periods of 0-8 hrs and 8-24 hours. The concentration of urea at the end of 8 hr period rose in the lumen compartment compared to the dialysate compartment, indicating an active uptake of urea by the probiotics in the lumen during the first phase of the experiment. Thereafter measurements made at the end of the 24-hr period revealed a reversal in the concentration gradient, where urea in the lumen was lower than that of the dialysate compartment. This indicates that the urea accumulated in the lumen during the first phase was actively metabolised by the probiotics in the second phase (8–24-time window) of the experiment. However, both runs of the experiment recorded a net deficit of urea compared to the total urea pumped into the system in 24 hrs, with run 1 and 2 recording total urea consumptions of 21.1 and 15.3 mmol respectively under the influence of 10^10 CFUs.

With regards to ammonium ion production, as expected the lumen had a higher local gradient of ammonium ion concentration than the dialysate compartment as the urea breakdown takes places only in the lumen compartment. Both experiments recorded a higher differential gradient at the end of 8hr period within the lumen compared to the dialysate compartment (Tables S3-S6). This higher gradient was also maintained in exp A at the end of 24hrs with 48.2 mM in the lumen versus 33.9 mM in the dialysate compartment in run 1. In both runs the molar ratio of total ammonia produced vs total urea consumed was found to be <1.0 (0.55 for exp A and 0.44 for exp B), indicating that a part of the ammonia produced as byproduct from urea metabolism was subsequently utilised by the probiotics as a nitrogen source (Fig 5). Taken collectively, the results show that the tested probiotics were capable of not only breaking down urea but also utilizing ammonia as a nitrogen source.

**Fig 5.**
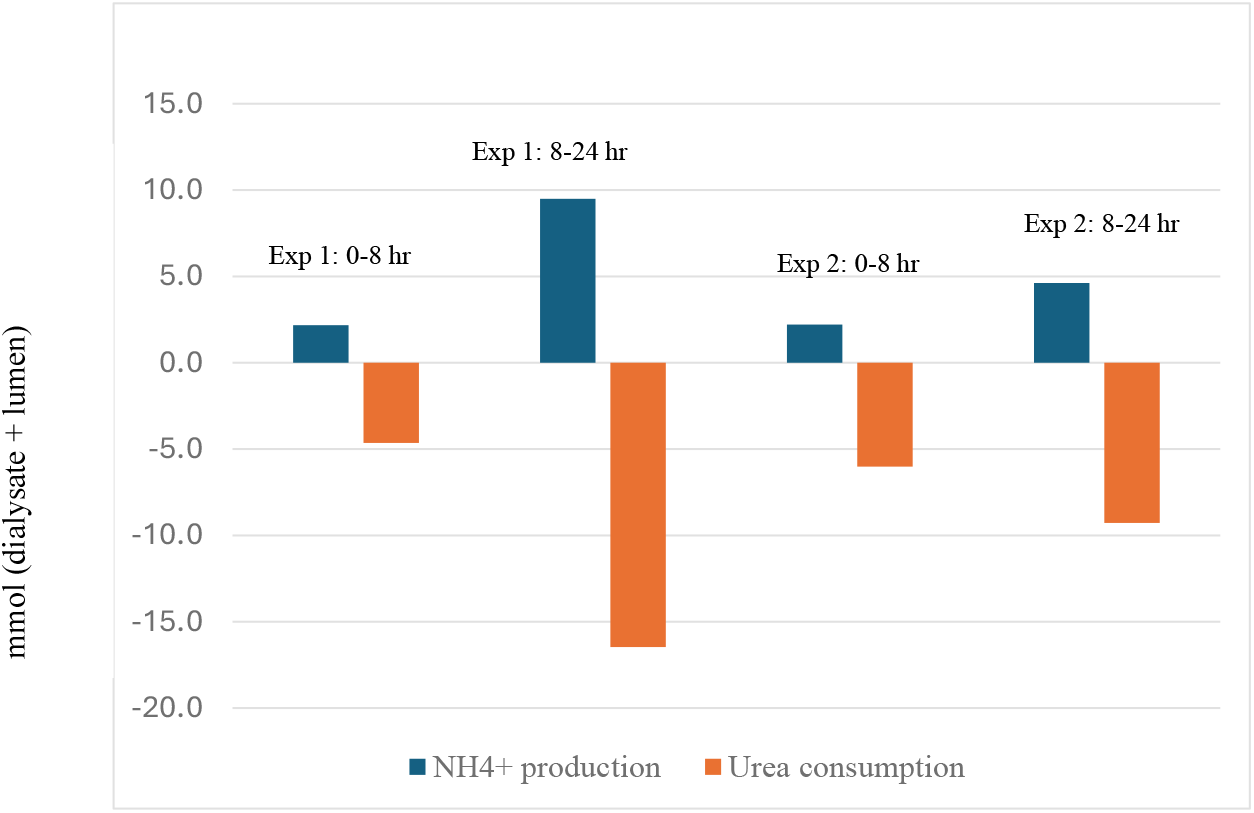
Urea breakdown and ammonium ion production in two individual runs of the TIM-2 system. In both runs, the molar ratio of total ammonia produced vs total urea consumed was found to be <1.0 (0.55 for exp A and 0.44 for exp B), indicating that a part of the ammonia produced as byproduct from urea metabolism was subsequently utilised by the probiotics as a nitrogen source. Taken collectively, the graph shows that the tested probiotics were capable of not only breaking down urea but also utilizing ammonia as a nitrogen source.

This study aimed to understand the effects of a probiotic mixture on urea metabolism in the TIM-2 colon model. The dialysis membrane in the colon compartment allows for the continuous removal of metabolites, preventing their accumulation. The impact of the probiotic test product on urea conversion was determined by measuring urea and ammonia in the dialysate and lumen samples. Both experiments demonstrated the conversion of urea by the probiotic bacteria. This led to the production of ammonia, which likely was partially utilized by the probiotic mixture. From the start of the TIM-2 experiments (t=0) the bacterial density was approximately 10^12 in the colonic lumen, which is similar to the density targeted with fecal inoculation of the model and comparable to *in vivo* colonic bacterial densities. The bacterial counts were similar between the experiments with and without urea, demonstrating that the concentration of urea was not toxic for the bacteria. The lower bacterial counts at the start of the adaptation period (t=-16 hours) indicate that the overnight adaptation, before adding the urea, is required for optimal performance of the test product and thus this overnight adaptation was applied to all experiments. Using bead beating to lyse the cells did not increase the measured metabolite concentrations. Possibly the process without the use of lysis buffer was not sufficient to release all cell content. It is also possible that the freeze/thaw cycle itself already releases most of the bacterial metabolites, or no significant amount was stored inside the cells. In urea experiment (A) the highest amount of ammonia was detected in the lumen sample at 24 hours. This might indicate that the highest conversion of urea occurred between 8 and 24 hours.

## Discussion

The findings of this study provide compelling evidence for the efficacy of specific probiotic strains in reducing uremic toxins, including urea, p-cresol, and indole. Given the limitations of conventional dialysis in completely removing these toxins, the use of probiotics presents an innovative and complementary approach to mitigate their detrimental effects. The study observed that all probiotic strains demonstrated significant activity in reducing urea levels *in vitro*. Training probiotic bacteria on toxic media using a gradient plate methodology offers a strategic approach to isolate and enhance strains with high toxin-degradation efficiency. Further training in liquid media confirmed the toxin degradation potential of the probiotic strains identified on gradient plates, with significant variation in efficiency among different species. The oscillatory pattern observed in urea breakdown capacity warrants further investigation to understand the underlying mechanisms and optimize degradation efficiency.

Experiments in the *in vitro* gut model offered a more controlled environment for testing the efficacy of the probiotic strains. The TIM-2 is an advanced in vitro gut model designed to simulate the physiological conditions of the human gastrointestinal tract, making it a valuable tool for assessing the effects of probiotics. One of the key advantages of the TIM-2 system is its ability to replicate the dynamic and complex conditions of the human gut. It can simulate the acidic environment of the stomach, the enzymatic activity in the small intestine, and the fermentation processes occurring in the colon. These conditions are critical for understanding how probiotics survive, proliferate, and exert their effects in the human body. Further, the model’s ability to simulate human gut conditions reduces the necessity of in vivo studies and accelerates the research process. Our results indicate that the probiotics were able to survive the harsh conditions of the stomach and small intestine to reach the colon, where they exert their beneficial effects.

However, the limitation of the TIM-2 colon model study is that no sampling took place between 8 and 24 hours (overnight), which would allow a more precise window of maximum urea conversion to be established. In a potential follow-up study additional sampling points could be included. In addition it might be worthwhile considering a longer experimental set-up (e.g. 72 hours) to get a more in depth understanding of the cycles of urea conversion and the adaptation of the bacteria to urea exposure. Lastly, it would be of interest to study the clinical probiotic dose in the TIM-2 colon model inoculated with active human microbiota. The peak in ammonia concentration described for experiment (A) was not observed in urea experiment (B). It was hypothesized that the conversion cycles of both experiments were not synchronized and might vary in its duration. This difference possibly occurred due to variation in the upregulation of gene expression, which could be an area of further investigation (e.g. with the remaining lumen samples). It was also observed that the net urea consumption was higher than the ammonia production. This could indicate that part of the ammonia produced after urea breakdown was consumed by the probiotic bacteria and could prevent the potentially harmful accumulation of ammonia. It would be of interest to understand the mechanisms involved in this process in more detail. While our study demonstrated that the probiotics were capable of consuming ammonia to some extent, they were unable to completely eliminate it. Ammonia accumulation can exacerbate metabolic acidosis, disrupt nitrogen homeostasis, and contribute to neurotoxic effects such as cognitive impairment, encephalopathy, and confusion. Additionally, increased systemic ammonia levels have been linked to heightened inflammation and oxidative stress, both of which are significant contributors to the progression of kidney disease and cardiovascular complications. Furthermore, ammonia can lead to an increased production of ammonium, which, in turn, can accelerate the loss of bone mineral density by altering acid-base balance and promoting osteodystrophy. Given these deleterious effects, any therapeutic strategy that inadvertently elevates ammonia levels without adequately addressing its clearance poses a substantial risk to patients. Therefore, understanding the implications of ammonia accumulation and the importance of further research into mitigating its harmful effects is paramount for the successful translation of this approach into clinical application.

Our findings highlight both the potential and the challenges of probiotic-driven urea breakdown as a novel treatment strategy. The ability of the engineered probiotics to partially consume ammonia suggests a valuable secondary function, yet their inability to fully eliminate ammonia underscores a crucial limitation that must be addressed before clinical translation. This result emphasizes the need for further optimization, such as enhancing the probiotic strains to more efficiently metabolize ammonia, co-administering ammonia-scavenging agents, or developing synergistic microbial communities with complementary metabolic functions. Additionally, investigating alternative pathways to safely excrete or detoxify ammonia will be critical for ensuring patient safety and efficacy of this therapeutic approach. Addressing the challenge of ammonia accumulation could significantly improve the viability of probiotic-based interventions and potentially revolutionize metabolic management in ESKD patients. Given the global burden of kidney disease and the limited treatment options available beyond dialysis and transplantation, exploring innovative solutions such as engineered probiotics represents an urgent and impactful scientific endeavour. In conclusion, while our study demonstrates the potential of trained probiotics in reducing urea, indole and p-cresol levels, the associated ammonia production presents a significant challenge that warrants further investigation. Understanding and mitigating ammonia toxicity will be essential for advancing this promising therapeutic approach into a clinically viable treatment. Future research aimed at optimizing probiotic metabolism, incorporating additional detoxification mechanisms, and evaluating long-term safety and efficacy in human subjects will be vital to fully harnessing the potential of this strategy for improving patient outcomes.

## Author Contributions

All authors made a significant contribution to the work reported. These contributions included the conception, study design, execution, data acquisition, analysis and interpretation. All authors contributed to the creation of the manuscript that included, drafting, revising and final approval of the submitted manuscript. Lastly, all authors agreed on the desired journal to submit the work.

