## Supplemental Material for "Evaluation of Probiotic Bacteria for the Reduction of Urea, p-Cresol, and Indole Levels"

**Supplementary Methods**

**Gradient plate preparation:** Gradient plates were prepared using rectangular plates, with 2/3 of the plate divided into 10 zones for growth media and the remaining 1/3 labelled as Zone 11. Plates were positioned at an angle, and growth agar was poured until it reached the 10th zone. After ~80% solidification, the plates were placed horizontally, and MM2 or MM3 agar was poured to cover the surface completely, followed by full solidification. For culture preparation, ATCC strains were revived, sub-cultured, and glycerol stocks prepared. Cultures were incubated under the recommended ATCC conditions for temperature and oxygen. Post-incubation, isolated colonies were inoculated into growth medium for a second passage, followed by centrifugation at 4500 RPM. Cells were washed twice with MM1, and the optical density was adjusted to 0.4 at 600 nm to serve as inoculum (~1 CFU/µL). A 100 µL inoculum was spread evenly on gradient plates and incubated under appropriate conditions. Controls were established by streaking the inoculum on growth media for growth control and using MM1 and agar media for sterility controls. For toxin screening, gradient plates were imaged post-incubation to identify bacterial growth in the highest toxin-concentration zones. Colonies from these zones were picked, spread onto Zone 11, and incubated further. This process was repeated until growth reached the highest toxin concentration or growth arrest was observed after 48 hours. Images were captured at 12-hour intervals to confirm growth arrest. Mutants were separated by re-streaking on fresh agar plates with the same toxin and nutrient concentrations as the original isolation zones. Each mutant strain was subsequently cultured in broth media, maintaining appropriate toxin and nutrient concentrations, and glycerol stocks were prepared.

To optimize bacterial inoculum for obtaining isolated colonies on agar plates, the initial inoculum was standardized using optical density measurements at 600 nm, followed by serial dilution and plating. Cell suspensions were prepared from overnight broth cultures, with the OD adjusted to 0.4–0.6 at 600 nm. Bacterial suspensions were then diluted to achieve a desired viable cell concentration of 50–100 CFU per 100 µL. Lactobacillus rhamnosus and Bifidobacterium longum were used as inoculum strains. For log-phase culture optimization, glycerol stock cultures were subcultured into broth medium and incubated overnight under the appropriate growth conditions. A volume of 100–200 µL from the overnight culture was transferred into 10 mL of sterile growth medium and incubated at 37°C for 6–8 hours in a shaker at 150–200 RPM. OD was measured at 600 nm, and serial dilutions were performed in MM1. Plating was conducted on the respective agar medium, followed by incubation under appropriate growth conditions. Colony-forming unit counts were recorded for each strain to evaluate growth

**Probiotic strains and growth conditions:** The incubation conditions and growth media for various microbial strains (and their ATCC codes) were as follows: Bacillus pasteurii (11859) was incubated at 30°C for 48 to 72 hours under aerobic conditions using Bacillus pasteurii NH4-YE medium. Bacillus coagulans (7050), Erwinia herbicola (33243), Streptococcus faecium (BAA-2320), and Streptococcus thermophilus (19258) were incubated at 37°C for 24 hours under aerobic conditions using nutrient agar or nutrient broth, or Brain Heart Infusion Agar/Broth, depending on the strain. Bifidobacterium longum (15707), Bifidobacterium breve (15700), and Bifidobacterium infantis (15697) were incubated at 37°C for 24 to 48 hours or 48 to 72 hours under anaerobic conditions with Modified Reinforced Clostridial medium. Lactobacillus casei (15008), Lactobacillus bulgaricus (11842), Lactobacillus rhamnosus (21052), Lactobacillus plantarum (BAA-793), and Lactobacillus gasseri (33323) were incubated at 37°C for 24 to 48 hours under aerobic conditions with 5% CO2, using Lactobacilli MRS Agar or Broth. Lactobacillus acidophilus (BAA-2845) was incubated at 37°C for 24 hours under anaerobic conditions with Lactobacilli MRS Agar or Broth. Akkermansia muciniphila (BAA-835) was incubated at 37°C with shaking for 2 to 7 days under anaerobic conditions with 100% N2, using Brain Heart Infusion Agar/Broth. These conditions ensured optimal growth and maintenance of each microbial strain.

**Urea/Ammonia Determination:** The urea breakdown assay was performed to measure total urea degradation in media and quantify the ammonia released using the Nessler’s reagent method. A standard curve prepared with ammonium chloride a maximum concentration of 100 µg/mL and serially diluted two fold to obtain a 10-fold dilution series. For each standard, 1 mL of solution was supplemented with 20 µL of sodium potassium tartrate solution and mixed thoroughly. Then, 30 µL of Nessler’s reagent was added, and the mixture was incubated at 25°C for 10 minutes to allow color development, during which any precipitate formation was observed. Absorbance was measured at 420 nm using water as a blank. A standard curve was generated by plotting the ammonia concentration (µg/mL) on the x-axis and the corresponding optical density values at 420 nm on the y-axis. For test samples, 1 mL of the media was supplemented with 20 µL of sodium potassium tartrate solution, followed by 30 µL of Nessler’s reagent. The mixture was incubated at 25°C for 10 minutes, and any precipitate formation was noted. The absorbance was then measured at 420 nm, using water as a blank. If the OD exceeded the linear range of the standard curve, the samples were diluted to 1:10 or 1:50 prior to analysis. Ammonia concentrations in the samples were calculated by comparing the OD values to the standard curve. Negative controls containing media without urea were included to account for background ammonia levels.

**LCMS measurement of p-cresol and indole:** The quantification of p-cresol and indole in test samples was performed using liquid chromatography-mass spectrometry. For sample preparation, 4 µL of aqueous calibration control (CC) and quality control (QC) solutions were mixed with 96 µL of blank growth media and vortexed for uniformity. Test samples were diluted based on their target compound: indole was diluted 2-fold, and p-cresol was diluted 5-fold by adding 4 µL of the test sample to 16 µL of blank growth media and vortexing. For indole analysis, 50 µL of the prepared CC, QC, and test sample solutions were extracted with 200 µL of acetonitrile containing internal standards (telmisartan and tolbutamide, 100 ng/mL each). For p-cresol analysis, 20 µL of the prepared CC, QC, and test sample solutions were similarly extracted with 200 µL of the same acetonitrile mixture. All extracted solutions were vortexed at 1000 rpm for 10 minutes, centrifuged at 4000 rpm for 10 minutes, and 130 µL of the supernatant was transferred to a 96-well plate for LC-MS/MS analysis.

Chromatographic analysis was performed using a Shimadzu Nexera X2 HPLC system coupled to a SCIEX Triple Quad 4500 mass spectrometer with Analyst 1.7.0 software. Separation was achieved using a Kinetex C18 column (100 × 3.0 mm, 5 µm), and ionization was conducted using a heated nebulizer source. Bacterial strains were grown in 12.5% pooled growth media and 87.5% MM1 with or without 20% dextrose, supplemented with 0.5 mM toxin (indole or p-cresol), and analyzed under the specified conditions. Calibration curves for indole (0.4–400 µM) and p-cresol (2–2000 µM) were prepared prior to the sample run, and QC samples were analyzed twice, before and after sample injections, with all results falling within the linear range. Samples with concentrations exceeding the calibration range were diluted appropriately. For indole, the control group showed concentrations of 386 µM and 421 µM (with 20% dextrose), while the test group showed 258 µM and 216 µM, corresponding to reductions of 127.5 µM in MM1 and 205 µM in MM1 + 20% dextrose conditions. For p-cresol, all samples showed concentrations exceeding 0.5 mM, with no significant differences observed between control and test groups. Data analysis confirmed that all sample concentrations fell within the established calibration range for their respective toxins.

**Invitro Gut Model:** The TNO invitro gut model (TIM-2 system) was employed in this study, which is elaborated in detail elsewhere^16^. In the TIM-2 System, the ascending, transverse and descending colon conditions was simulated, including gradual pH increase from 5.5 towards 7.0 and decrease of fermentable carbohydrates. The system was kept under strict anaerobic conditions. The TIM-2 System was inoculated with a mixture of equal ratios of probiotic strains with a cumulative 10 billion CFUs in 125 ml in simulated ileal effluent medium(SIEM) and dialysis solution. The SIEM simulates the material reaching the colon (Western diet) and it was used as standard feeding, denoted as control^17^. It contains indigestible carbohydrates (pectin, xylan, arabinogalactan, amylopectin and starch), protein, vitamins, and bile^17^. Tween 80 was omitted because it previously was found to hinder analysis of carbohydrate degradation products^16^. The pH was adjusted to 5.8 to simulate the pH from the proximal colon. Dialysis solution contained (per litre): 2.5 g K2HPO4·3H2O, 4.5 g NaCl, 0.005 g FeSO4·7H2O, 0.5 g MgSO4·7H2O, 0.45 g CaCl2·2H2O, 0.05 g bile and 0.4 g cysteine∙HCl, plus 1 mL of the vitamin mixture; pH 5.8 (Aguirre et al., 2015). All medium components were purchased at Tritium Microbiology (Eindhoven, the Netherlands).

One pilot TIM-2 run was performed with the probiotic mixture and urea to evaluate and finalize the experimental set-up, before conducting the main study. The main study included two runs with urea and one control run without urea. Each run lasted 24 hours. The TIM-2 units was flushed with nitrogen prior to inoculation and during the entire experiment. At the start of the adaptation period, the TIM-2 System was inoculated with approximately 35 ml of thawed probiotic mixture and 90 ml dialysis fluid. The probiotic mixture was allowed to adapt to the model conditions and SIEM for approximately 16 hours at pH 5.5. After the adaptation period, the 24 hours test period started. At this timepoint (t=0) filter-sterilized urea was added to the colonic lumen to reach a concentration of 25mM (1.5 g/L), mimicking the levels found in renal patients. As low molecular weight compounds are continuously removed from the lumen via dialysis, urea was continuously added at a fixed concentration of 25mM (1.5 g/L) in the dialysis liquid. The speed of the dialysis liquid is 1.5 ml/min. This results in the addition of approximately 2.25 mmol urea in 90mL per hour. The outcoming dialysis volumes were recorded. Subsequently, samples were collected (referred to as t0h). Samples from lumen and dialysate out were collected every ∼24 h (t0h, t24h, t48h and t73h). The collected samples were subsequently analysed for urea and ammonia levels.

**Sample processing from in vitro gut model:** During passage of the probiotic mixture through the TIM-2, the following samples were collected: Dialysate, corresponding to the bio-accessible fraction (Figure S1). This represents the amount of metabolites available for absorption from the colon. Lumen sample, corresponding to the colonic material (Figure S1b, part G). For the metabolite analysis, dialysate and lumen samples were collected at: t = 0, 8 and 24 hours after the start of the test period. The lumen samples in the main phase were exposed to 10 cycles of 1 minute bead beating using zirconium beads, to lyse the bacterial cells present. This resulted in a total of 24 samples (4 TIM-2 runs x 3 sampling time points x 2 types of samples – lumen and dialysate) analyzed for urea and ammonia. In addition, samples at these points were stored for potential future UPLC analysis. Luminal samples were collected at t = 0, 8 and 24 hours to determine the bacterial density of the bacterial strains in the probiotic mixture.

| **Indole** | | | |
| --- | --- | --- | --- |
| **Sample Name** | **Nominal Conc. (mM)** | **Calculated Conc. (mM)** | **Accuracy (%)** |
| BLANK | 0.00 | N/A | N/A |
| BLANK+IS | 0.00 | N/A | N/A |
| Indole CC-1 | 0.51 | 0.52 | 101.04 |
| Indole CC-2 | 1.02 | 1.01 | 99.31 |
| Indole CC-3 | 2.05 | 2.00 | 97.37 |
| Indole CC-4 | 4.10 | 4.13 | 100.67 |
| Indole CC-5 | 6.30 | 5.79 | 91.9 |
| Indole CC-6 | 9.00 | 9.62 | 106.93 |
| Indole CC-7 | 12.00 | 12.64 | 105.3 |
| Indole CC-8 | 16.00 | 15.48 | 96.76 |
| Indole CC-9 | 20.00 | 20.14 | 100.72 |
| LQC-1 | 1.54 | 1.48 | 96.42 |
| MQC-1 | 10.00 | 10.05 | 100.51 |
| HQC-1 | 18.00 | 16.88 | 93.75 |
| LQC-2 | 1.54 | 1.58 | 102.85 |
| MQC-2 | 10.00 | 9.98 | 99.78 |
| HQC-2 | 18.00 | 16.63 | 92.40 |
| Linearity range: 0.51-20 mM | | | |

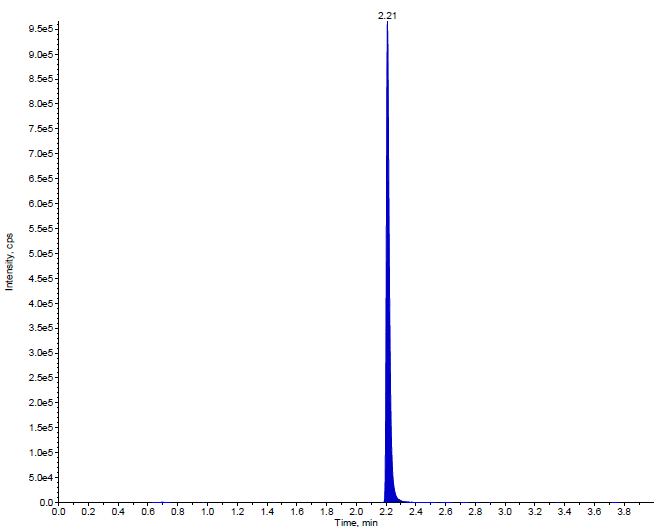

**Indole Peak**

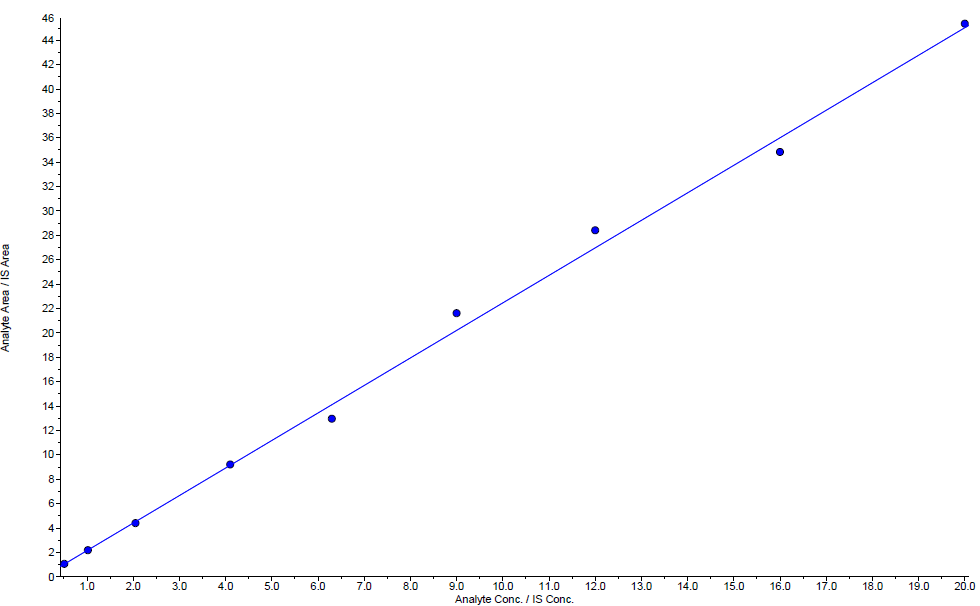

**Indole Calibration Curve**

CC: Calibration curve standard

LQC: Lower quality control

MQC: Middle quality control

HQC: Higher quality control

Quality controls samples are run bracketing the test samples and batch is accepted if quality controls are within in-house acceptance limit (80-120%).

**Fig S1. LC-MS calibration data for indole with a established linearity range of 0.5 to 20 mM.**

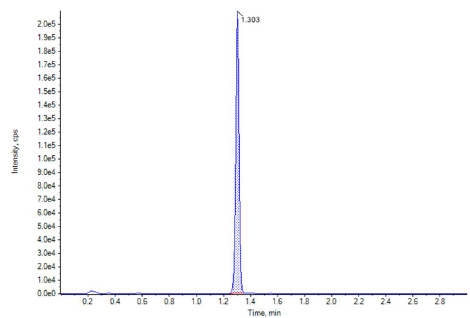

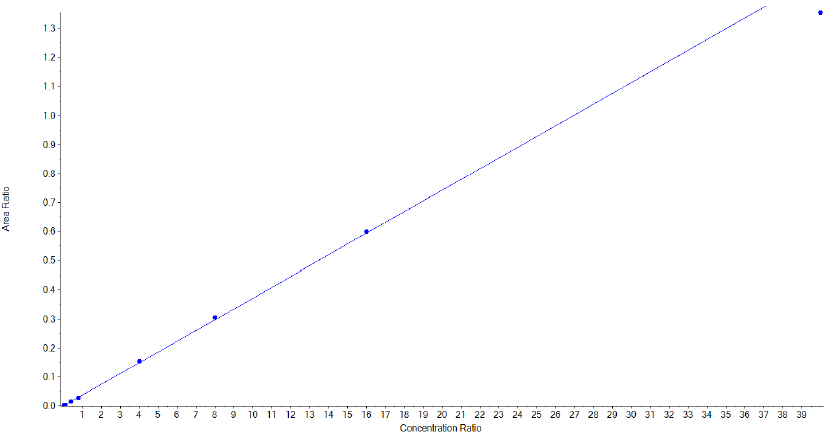

**p-cresol Calibration Curve**

**p-cresol Peak**

| **p-cresol** | | | |
| --- | --- | --- | --- |
| **Sample Name** | **Nominal Conc. (mM)** | **Calculated Conc. (mM)** | **Accuracy (%)** |
| **BLANK** | **0.00** | **N/A** | **N/A** |
| **BLANK+IS** | **0.00** | **N/A** | **N/A** |
| **p-cresol CC-1** | **0.51** | **0.50** | **98.94** |
| **p-cresol CC-2** | **1.02** | **1.04** | **101.51** |
| **p-cresol CC-3** | **2.05** | **2.12** | **103.33** |
| **p-cresol CC-4** | **4.10** | **3.90** | **95.11** |
| **p-cresol CC-5** | **6.30** | **6.16** | **97.73** |
| **p-cresol CC-6** | **9.00** | **9.43** | **104.80** |
| **p-cresol CC-7** | **12.00** | **12.23** | **101.93** |
| **p-cresol CC-8** | **16.00** | **15.88** | **99.25** |
| **p-cresol CC-9** | **20.00** | **19.48** | **97.39** |
| **LQC-1** | **1.54** | **1.27** | **82.67** |
| **MQC-1** | **15.00** | **12.97** | **86.47** |
| **HQC-1** | **18.00** | **16.31** | **90.60** |
| **LQC-2** | **1.54** | **1.20** | **78.20** |
| **MQC-2** | **15.00** | **12.48** | **83.23** |
| **HQC-2** | **18.00** | **15.61** | **86.74** |
| **Linearity range: 0.51-20 mM** | | | |

CC: Calibration curve standard

LQC: Lower quality control

MQC: Middle quality control

HQC: Higher quality control

Quality controls samples are run bracketing the test samples and batch is accepted if quality controls are within in-house acceptance limit (80-120%).

**Fig S2. LC-MS calibration data for indole with a established linearity range of 0.5 to 20 mM.**

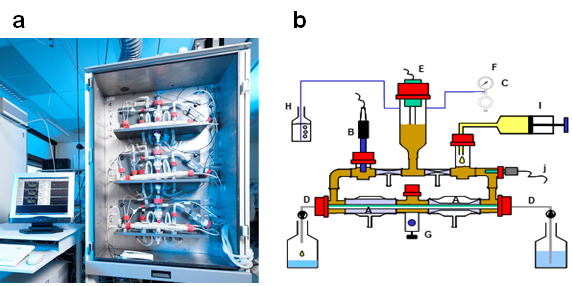

**Fig S3**. **a)** TIM-2 System. **b)** Schematic overview of the TIM-2 System. A. mixing units; B. pH electrode; C. alkali pump; D. dialysis liquid circuit with hollow fibers; E. level-sensor; F. N_2_ gas inlet; G. sampling port; H. gas outlet; I. 'ileum effluent' container; J. temperature sensor.

**Table S1.** Ammonia quantification by Nessler’s reagent

| **Ammonia Quantification (NH_4_Cl (μg/ml)** | | | | | | |
| --- | --- | --- | --- | --- | --- | --- |
| **ATCC strains** | **Day 1** | **Day 2** | **Day 3** | **Day 4** | **Day 5** | **Day 6** |
| L rhamnosus (21052) | NA | NA | NA | 5793 | 3643 | 4423 |
| L plantarum (BAA-793) | 279 | 2030 | 2267 | 5030 | 2428 | 4119 |
| S thermophilus (19258) | 983 | 2043 | 3291 | 4846 | 2770 | 4035 |
| S faecium (BAA-2320) | 249 | 2268 | 1380 | 2159 | 3017 | 4437 |
| B longum (15707) | 1231 | 4532 | 6244 | 7276 | 5937 | 5446 |
| B breve (15700) | 900 | 1798 | 2358 | 3445 | 2755 | 3953 |
| L casei (15008) | 244 | 2130 | 2477 | 4914 | 3044 | 3368 |
| L acidophilus (BAA-2845) | 2476 | 4209 | 2660 | 4738 | 6281 | 5692 |
| L bulgaricus (11842) | 228 | 1631 | 2987 | 6102 | 2768 | 4158 |
| L gasseri (33323) | 2351 | 4853 | 3251 | 4305 | 6302 | 4620 |
| A muciniphila (BAA-835) | 28 | 3132 | 5714 | 3779 | 4387 | 3133 |
| B infantis (15697) | 314 | 2949 | 2013 | 2652 | 3357 | 4753 |
| B coagulans (7050) | 1027 | 1524 | 3399 | 4938 | 3670 | 5740 |
| E herbicola (33243) | 23 | 278 | 198 | 1069 | 1614 | 2680 |
| B pasteurii (11859) | 96 | 1113 | 1630 | 4032 | 2058 | 2429 |
| B subtilis (6633) | 872 | 3498 | 3138 | 7759 | 3729 | 3756 |

**Table S2.** OD_600_ values of probiotic strains across 6 days

| **ATCC strains** | **Day 1** | **Day 2** | **Day 3** | **Day 4** | **Day 5** | **Day 6** |
| --- | --- | --- | --- | --- | --- | --- |
| L rhamnosus (21052) | NA | NA | NA | 0.165 | 0.171 | 0.166 |
| L plantarum (BAA-793) | 0.154 | 0.140 | 0.146 | 0.138 | 0.126 | 0.135 |
| S thermophilus (19258) | 0.199 | 0.311 | 0.138 | 0.280 | 0.270 | 0.268 |
| S faecium (BAA-2320) | NA | NA | 0.162 | 0.150 | 0.138 | 0.130 |
| B longum (15707) | 0.192 | 0.270 | 0.259 | 0.253 | 0.182 | 0.228 |
| B breve (15700) | 0.121 | 0.109 | 0.190 | 0.281 | 0.156 | 0.142 |
| L casei (15008) | 0.158 | 0.155 | 0.141 | 0.139 | 0.135 | 0.142 |
| L acidophilus (BAA-2845) | 0.335 | 0.300 | 0.322 | 0.346 | 0.314 | 0.286 |
| L bulgaricus (11842) | 0.168 | 0.169 | 0.150 | 0.148 | 0.142 | 0.146 |
| L gasseri (33323) | 0.322 | 0.386 | 0.362 | 0.348 | 0.336 | 0.329 |
| A muciniphila (BAA-835) | 0.221 | 0.198 | 0.190 | 0.179 | 0.293 | 0.163 |
| B infantis (15697) | NA | NA | 0.196 | 0.192 | 0.185 | 0.179 |
| B coagulans (7050) | 0.161 | 0.275 | 0.244 | 0.235 | 0.223 | 0.215 |
| E herbicola (33243) | NA | NA | 0.152 | 0.169 | 0.198 | 0.203 |
| B pasteurii (11859) | 0.189 | 0.225 | 0.316 | 0.283 | 0.256 | 0.243 |
| B subtilis (6633) | 0.351 | 0.351 | 0.354 | 0.339 | 0.337 | 0.468 |

**Table S1.** Conditions and parameters simulated in TIM-2

| **Simulated conditions** | **Parameters** |
| --- | --- |
| Substrate (t=0-8h) | SIEM with carbohydrates |
| Substrate (t=8-24h) | SIEM without carbohydrates |
| pH ascending colon (t = 0-8h) | 5.5 - 6.5 |
| pH transverse colon (t = 8-16h) | 6.5 - 7.0 |
| pH descending colon (t = 16-24h) | 7.0 |
| Residence time in TIM-2 during test period | 24 hours |

**Table S2.** Concentration and amount of ammonia at different time intervals in the dialysate and time points in the lumen samples. The absolute amounts of ammonia are calculated taking into account the volumes of the dialysate and lumen respectively.

|  |  | **free** | **absolute** | **dialysate+lumen** | **rate** |
| --- | --- | --- | --- | --- | --- |
|  |  | **[NH4+]** | **[NH4+]** | **[NH4+]** | **[NH4+]** |
|  | **Time (h)** | **(mM)** | **mmol** | **mmol** | **mmol/hr/total CFU** |
| **Dialysate** | **-16-0** | 0.6 | 0.9 |  |  |
|  | **0-8** | 1.2 | 0.8 | 1.0 | 0.12 |
|  | **8-24** | 4.2 | 6.2 | 9.7 | 0.61 |
| **Lumen** | **0** | 1.7 | 0.2 |  |  |
|  | **8** | 2.5 | 0.3 |  |  |
|  | **24** | 30.5 | 3.8 |  |  |

**Table S3.** Concentration and amount of urea at different time intervals in the dialysate and time points in the lumen samples. The absolute amounts of ammonia are calculated taking into account the volumes of the dialysate and lumen respectively.

|  |  | **total** |  | **absolute** | **dialysate+lumen*** | **rate** |
| --- | --- | --- | --- | --- | --- | --- |
|  |  | **[NH4+]** | **[urea]** | **[urea]** | **[urea]** | **[urea]** |
|  | **Time (h)** | **(mM)** | **(mM)** | **mmol** | **mmol** | **mmol/hr/total CFU** |
| **Dialysate** | **-16-0** | 0.9 | 0.1 | 0.2 |  |  |
|  | **0-8** | 40.1 | 19.4 | 14.1 | -1.4 | -0.17 |
|  | **8-24** | 51.2 | 23.5 | 35.1 | -6.1 | -0.38 |
| **Lumen** | **0** | 3.4 | 0.9 | 0.1 |  |  |
|  | **8** | 93.3 | 45.4 | 5.7 |  |  |
|  | **24** | 38.3 | 3.9 | 0.5 |  |  |

**Table S4.** Concentration (mM) and amount (mmol) of ammonia at different time intervals in the dialysate and time points in the lumen samples for the experiments with urea (n=2) and a blank experiment. The production of ammonia was calculated taking into account the volumes of the dialysate and lumen respectively.

|  |  |  | **free** | **absolute** | **dialysate+lumen** | **rate** |
| --- | --- | --- | --- | --- | --- | --- |
|  |  |  | **[NH4+]** | **[NH4+]** | **[NH4+]** | **[NH4+]** |
|  |  | **Time (h)** | **(mM)** | **mmol** | **mmol** | **mmol/hr/total CFU** |
| **Blank** | **Dialysate** | **-16-0** | 0.8 | 1.2 |  |  |
|  |  | **0-8** | 1.6 | 1.2 | 1.4 | 0.17 |
|  |  | **8-24** | 1.5 | 2.3 | 2.0 | 0.12 |
|  | **Lumen** | **0** | 2.8 | 0.4 |  |  |
|  |  | **8** | 4.1 | 0.5 |  |  |
|  |  | **24** | 1.7 | 0.2 |  |  |
| **Urea** | **Dialysate** | **-16-0** | 0.8 | 1.3 |  |  |
| **(A)** |  | **0-8** | 2.8 | 2.1 | 2.2 | 0.27 |
|  |  | **8-24** | 3.0 | 4.4 | 9.5 | 0.59 |
|  | **Lumen** | **0** | 3.2 | 0.4 |  |  |
|  |  | **8** | 3.9 | 0.5 |  |  |
|  |  | **24** | 44.8 | 5.6 |  |  |
| **Urea** | **Dialysate** | **-16-0** | 0.9 | 1.3 |  |  |
| **(B)** |  | **0-8** | 2.9 | 2.1 | 2.2 | 0.28 |
|  |  | **8-24** | 3.2 | 4.7 | 4.6 | 0.29 |
|  | **Lumen** | **0** | 2.9 | 0.4 |  |  |
|  |  | **8** | 3.6 | 0.5 |  |  |
|  |  | **24** | 3.3 | 0.4 |  |  |

**Table S5.** Concentration (mM) and amount (mmol) of urea at different time intervals in the dialysate and time points in the lumen samples for the experiments with urea (n=2) and a blank experiment. The absolute amounts of urea formed were calculated taking into account the volumes of the dialysate and lumen respectively.

|  |  |  | **total** |  | **absolute** | **dialysate+lumen*** | **rate** |
| --- | --- | --- | --- | --- | --- | --- | --- |
|  |  |  | **[NH4+]** | **[urea]** | **[urea]** | **[urea]** | **[urea]** |
|  |  | **Time (h)** | **(mM)** | **(mM)** | **mmol** | **mmol** | **mmol/hr/total CFU** |
| **Blank** | **Dialysate** | **-16-0** | 1.0 | 0.1 | 0.2 |  |  |
|  |  | **0-8** | 1.6 | 0.0 | 0.0 | 0.1 | 0.02 |
|  |  | **8-24** | 1.5 | 0.0 | 0.0 | 0.2 | 0.01 |
|  | **Lumen** | **0** | 4.8 | 1.0 | 0.1 |  |  |
|  |  | **8** | 6.4 | 1.2 | 0.1 |  |  |
|  |  | **24** | 4.7 | 1.5 | 0.2 |  |  |
| **Urea** | **Dialysate** | **-16-0** | 1.0 | 0.1 | 0.1 |  |  |
| **(A)** |  | **0-8** | 37.0 | 17.1 | 12.9 | -4.6 | -0.58 |
|  |  | **8-24** | 33.9 | 15.5 | 22.9 | -16.5 | -1.03 |
|  | **Lumen** | **0** | 5.1 | 1.0 | 0.1 |  |  |
|  |  | **8** | 61.4 | 28.8 | 3.6 |  |  |
|  |  | **24** | 48.2 | 1.7 | 0.2 |  |  |
| **Urea** | **Dialysate** | **-16-0** | 1.1 | 0.1 | 0.1 |  |  |
| **(B)** |  | **0-8** | 32.0 | 14.6 | 10.6 | -6.0 | -0.75 |
|  |  | **8-24** | 42.8 | 19.8 | 29.3 | -9.3 | -0.58 |
|  | **Lumen** | **0** | 4.5 | 0.8 | 0.1 |  |  |
|  |  | **8** | 76.4 | 36.4 | 4.5 |  |  |
|  |  | **24** | 35.6 | 16.2 | 2.0 |  |  |

* corrected for urea addition over time
